# Gene family evolution in brassicaceous-feeding insects: Implications for adaptation and host plant range

**DOI:** 10.1101/2023.06.09.544424

**Authors:** Nitin Ravikanthachari, Carol L Boggs

**Author notes:** <u>Corresponding author:</u> Nitin Ravikanthachari. Division of Biological Sciences, University of Montana, Missoula, MT 59812, USA.

## Abstract

Herbivores have a defined range of hostplants that they can feed on, which is mediated by underlying detoxification and sensory repertoires. Insects that feed on Brassicaceae represent one of the striking examples of co-evolutionary arms race. Insects specialized on Brassicaceae have evolved specific mechanisms to detoxify mustard oils (glucosinolates), while generalist species use detoxification enzymes that act on a variety of substrates. Understanding the gene evolution of detoxification and sensory repertoire in specialist and generalist Brassicaceae feeders will shed light on the processes involved in mediating hostplant ranges in herbivores. We use a comparative phylogenomic approach in 12 lepidopterans that feed on Brassicaceae, ranging from specialist to pests in their host range to examine the gene family expansion of detoxification and sensory gene families. We found that gene family expansions and contractions were larger in generalist herbivores compared to specialist herbivores. Gene evolutionary rate of detoxification genes reflected hostplant range where generalists had a higher evolutionary rate of detoxification genes that act on wide substrates while specialists had a higher evolutionary rate in genes that conjugate toxic compounds to hydrophilic byproducts. Our analysis on the nitrile specifier gene, a key innovation for feeding on Brassicaceae, indicated pervasive purifying selection with lineage specific differences in selection. Our results add to the growing body of work addressing gene family evolution and its role in hostplant range and specialization in insects.

## Introduction

Co-evolutionary interaction between plants and their herbivores is often proposed as one of the major drivers of eukaryotic biodiversity on earth (Ehrlich & Raven, 1964; Futuyma & Agrawal, 2009; Wiens et al., 2015). The co-evolutionary arms race is a product of ongoing key adaptive innovations in chemical and physical defense in plants and behavioral and chemical adaptation to circumvent those defenses in herbivores (M. Berenbaum, 1983; M. R. Berenbaum et al., 1986; Edger et al., 2015; Wheat et al., 2007). As a result, many herbivores have evolved specific mechanisms to detoxify a narrow range of hostplant chemicals and thus feed and develop on them (Hardy & Otto, 2014; Jaenike, 1990; Joshi & Thompson, 1995). The hostplant range of an herbivore is thus dependent on multiple factors including the degree similarity of chemical defenses among host plants, evolutionary history of hostplant use, genetic variation in detoxification enzymes and diversity of sensory repertoire in herbivores (Awmack & Leather, 2002; Bernays & Graham, 1988; Gripenberg et al., 2010).

Insect detoxification systems involve three distinct phases, which help in converting toxic plant chemicals to inert substances that are then excreted (Heidel-Fischer & Vogel, 2015; Kant et al., 2015; Rane et al., 2019). Phase I detoxification consists of cytochrome p450 monooxygenases (P450s or CYPs) and carboxylesterases (COesterase) which introduce reactive and polar groups into plant toxic compounds by reduction or hydrolysis (Dermauw et al., 2020; Feyereisen, 1999; Ranson et al., 2002). Phase II detoxification enzymes which include Glutathione-S-Transferases (GSTs), UDP-glucosyltransferase (UDPGTs or UGTs) and Sulfotransferases add either glutathione, glucosyl or sulfate groups respectively to the byproducts from phase I or to toxic compounds themselves to increase their hydrophilicity (Cermak, 2008). Finally, enzymes in phase III such as ABC (ATP-binding cassettes) transporters that export these bound toxins to the extracellular matrix for excretion (Dermauw & Van Leeuwen, 2014).

Polyphagous herbivores (those that feed on a wide range of plant families with diverse chemical composition) and specialist herbivores (those that are specialized to feed on one plant family with a particular class of chemical profile) have different mechanisms by which they detoxify plant compounds. Polyphagous herbivores are known to have higher gene expression and increased gene copies of phase I and phase II general detoxification enzymes whereas specialist herbivores have evolved specific detoxification enzymes in addition to the general detoxification machinery (Calla et al., 2017; Edger et al., 2015; Wheat et al., 2007).

The interaction between the plant family Brassicaceae (mustard family) and its herbivores has been intensively studied in order to dissect the mechanisms underlying detoxification of specialized toxic compounds (Edger et al., 2015; Ratzka et al., 2002; Wheat et al., 2007). Brassicaceae have evolved glucosinolates (GLS), a class of secondary compounds that are stored separately from myrosinases (Wittstock & Halkier, 2002). Upon damage by herbivores, glucosinolates and myrosinases interact, followed by the hydrolysis of glucosinolates into highly toxic isothiocyanates (Halkier & Gershenzon, 2006). Brassicaceae are well defended as most herbivores are unable to process isothiocyanates. However, specialized herbivores including some species of Lepidoptera such as *Pieris sps.* (Pieridae) and *Plutella xylostella* (Plutellidae) have evolved mechanisms to redirect the GLS-myrosinase reaction to instead form nitriles that are less toxic (Heidel-Fischer & Vogel, 2015). *Pieris sps.* have evolved nitrile specifier protein (NSP), which is expressed in the midgut to detoxify GLS (Chew, 1977, 1980; Kuchernig et al., 2012). NSPs belong to the family of Insect allergen repeat, which is present in all insects and whose function is largely unknown. However, in *Pieris spp.*, evolution of NSP was a key innovation in the diversification of the genus as well as their ability to utilize Brassicaceae (Edger et al., 2015; Pomés et al., 1998; Wheat et al., 2007).

Apart from the above-mentioned specialist Lepidoptera, several species in the family Noctuidae (Lepidoptera) are known to feed on Brassicaceae in addition to feeding on a large range of hostplants. Several of these species are also well-known pest species including *Helicoverpa armigera*, *Spodoptera exigua* and *Trichoplusia ni* (Cho et al., 2008). Although they do not possess specialized enzymes mentioned above, they are able to feed on a diverse set of hostplant families including Brassicaceae through differential expression and regulation of the general detoxification machinery.

In addition to detoxification, sensory perception plays a crucial role in the hostplant range of insect herbivores. Sensory receptors are usually fine-tuned to the secondary metabolites present in the hostplant (Engsontia et al., 2014). For example, ovipositing females can differentiate between hosts and non-hosts as well as assess the nutritional quality of the hostplant using gustatory and olfactory receptors (Ozaki et al., 2011; Ryuda et al., 2013). In fact, the diversity of sensory receptors is positively correlated with hostplant range in insects (Engsontia et al., 2014). Sensory receptors consist of three classes: The first class of receptors are olfactory receptors that aid in recognition of hostplants and conspecifics (Brand et al., 2018; Missbach et al., 2014). Gustatory receptors are involved in taste perception and consists of various subclasses that can identify sweet, bitter, and salt compounds (Nei et al., 2008; Sánchez-Gracia et al., 2009). The final class of sensory receptors are ionotropic receptors that are ligand gated channels that are crucial in the perception of secondary metabolites (Rytz et al., 2013).

Understanding the causal processes of hostplant specialization and the mechanisms through which generalist and specialist herbivores interact with well-defended plant toxins can shed light on the evolution and maintenance of detoxification machinery, which has important practical implications in agricultural management of pest species. Dynamic changes in gene family sizes, neofunctionalization and gene duplications in detoxification genes are hypothesized to be among the causal mechanism for the evolution of hostplant range in herbivorous insects (Dermauw et al., 2020; Dermauw & Van Leeuwen, 2014; Halon et al., 2015; Suzuki et al., 2018). Therefore, understanding the evolution of gene family size and selection on specialized detoxification genes can help shed light on the mechanisms through which polyphagous and specialist herbivores feed on well defended plant toxins.

We here use a comparative phylogenetic approach by using high quality genomes of lepidopteran insects that are both specialist and generalist feeders to test: a) if generalist herbivores have larger gene family changes in detoxification and sensory gene families compared to specialists; b) if detoxification and sensory gene families in generalist species evolve faster than specialists and c) if Nitrile Specifier Protein (NSP) was under purifying or diversifying selection.

## Methods

### Data collection and quality assessment

We downloaded genomes of Lepidoptera which feed on Brassicaceae (table S1) from NCBI (Sayers et al., 2021) and Ensemble LepBase (Challi et al., 2016). The final list of species, the corresponding accession numbers and date accesses are provided in the supplementary table S1. We designated a species as a specialist herbivore if it only fed on Brassicaceae family and as a generalist herbivore if it fed on other plant families in addition to Brassicaceae. We assessed genome quality based on BUSCO scores with the Lepidoptera gene set consisting of 5218 BUSCO in BUSCO v. 4 (Simão et al., 2015). Genomes that showed a high percentage of fragmented or missing BUSCOs were excluded from downstream analysis.

### Structural and functional annotation

We softmasked the repeats in our genomes using redmask (Girgis, 2015). Softmasked genomes were run through the BRAKER2 (Brůna et al., 2021) annotation pipeline with AUGUSTUS for gene predictions, with the Arthropoda gene set from OrthoDB (Kriventseva et al., 2019) serving as the reference proteins. We used gffread from the cufflinks pipeline (Trapnell et al., 2012) to extract protein and CDS sequences following gene prediction by BRAKER2. We retained only one isoform per gene when genes were represented by multiple isoforms using agat_sp_keep_longest_isoform_pl function in AGAT (Dainat et al., 2020). The final protein set was cleaned of illegal characters such as “*” and “.” and used for functional annotation with EggNOG (Cantalapiedra et al., 2021) and InterProScan (Jones et al., 2014). We used the Arthropoda dataset for functional annotation and realigned all the terms to their respective protein families using the Pfam (Mistry et al., 2021) databse for EggNOG annotation. For InterProScan, we used the -appl Pfam -goterms to get information about protein family annotations. We ran BUSCO again on the final protein set using the lepidopteran gene set to assess completeness of the proteome.

### Orthologs inference and phylogenetic tree inference

We used OrthoFinder (Emms & Kelly, 2019) to predict orthologous protein groups (OGs). An orthogroup is a set of genes that have descended from a single gene from the last common ancestor (LCA) in a clade of species. The filtered protein set was used as input for Orthofinder and was run with default settings to predict OGs.

We additionally used Orthofinder for constructing our species and gene phylogenetic tree. We used the -M msa option in Orthofinder for multiple sequence alignment using MAFFT (Katoh & Standley, 2013). We used the aligned single copy orthologs for building our species phylogenetic tree and the aligned sequences for each OG to construct the respective gene trees using IQTree (Nguyen et al., 2015). We used the default setting under IQTree for inferring species and gene trees. The root of the species tree was time calibrated to 87 Myr to reflect the divergence of the three families used in the analysis (Espeland et al., 2018).

### Gene family evolution

CAFE v. 5.0 (Mendes et al., 2020) was used to analyze gene family evolution under a phylogenetic framework. We used the phylogenetic hierarchical orthogroups from OrthoFinder analysis to retrieve gene counts per species as input for CAFE. We first filtered OGs whose gene counts were greater than 100 in any single species as recommend by the developers. We additionally filtered OGs that showed high variance across species as they can lead to biases in gene family evolution estimation as well as preventing convergence among replicates in the analysis.

Once our test analysis using the base parameters showed convergence, we calculated the error in gene assembly and annotation in our dataset using the -e option in CAFE. We subsequently used the estimated error in our subsequent analysis to account for genome annotation errors. We used 4 different data sets for our gene family evolution analysis. Our first dataset included OGs comprised of all gene families after filtering of high variance and high copy numbers as indicated above, hereby referred to as “all genes”. Second, we generated a filtered dataset consisting of eight gene families involved in detoxification of plant compounds (p450 and COesterase in Step I, UGT, GST and ST in Step II, ABC transporter in Step III and insect cuticle protein and trypsin related to general digestive process), hereby referred to as “detoxification genes”. Our third dataset included a filtered dataset consisting of three gene families involved in sensory recognition in Lepidoptera (gustatory, ionotropic, and olfactory genes), hereby referred to as “sensory genes”. Our final filtered data set consisted of individual gene families involved in detoxification process and sensory recognition referred to as “single gene family”.

We ran CAFE in two different modes. Our first run consisted of estimating a single rate of change (λ) for each of the 4 datasets to calculate a “baseline” λ for the entire tree. In our second run, we estimated λ separately for specialists, generalists and *Plutella xylostella*. We calculated a separate λ for *P. xylostella* for 2 reasons: a) *Plutella xylostella* was indicated as an outgroup in our tree, and therefore, including it with the rest of the specialists would bias our λ estimates for specialists and b) *P. xylostella* belongs to a different family compared to all the other specialist feeders (*Pieris sps.*, Pieridae) and generalist feeders (Noctuidae) in our dataset. We ran CAFE multiple times on all our datasets to ensure convergence among runs and used those estimates for determining λ for gene family evolution.

### Signatures of selection on NSP gene

We used the HyPhy suite (Pond et al., 2005) on the webserver Datamonkey (Weaver et al., 2018) to test signatures of selection on the NSP gene. We first used the multiple aligned sequence of the OG corresponding to NSP from the OrthoFinder analysis and converted it to aligned CDS sequences using Pal2Nal (Suyama et al., 2006) to use as input in HyPhy. We used FEL (fixed effects likelihood) to investigate whether individual sites in the NSP gene were subjected to pervasive selection. FEL uses a ML method to infer nonsynonymous (dN) and synonymous (dS) substitution rate for each site. We ran FEL with 1000 parametric replicates and used an asymptotic chi-squared test to assess significance. To test if selection on the NSP gene in the *Pieris spp.* (specialist and butterfly) clade was different compared to the rest (generalists and/or moths) of the phylogeny, we used contrast-FEL with branches corresponding to *Pieris spp.* as foreground selection and the rest of the branches as background selection. We assessed significant differences in dN/dS based on a q-value threshold of <0.2. Finally, we used MEME (Mixed Effects Model of Evolution) to test if sites in the NSP gene were subjected to episodic selection.

## Results

### Genome quality, gene family statistics and phylogeny

Our analysis of twelve Lepidoptera genomes included 4 specialist Brassicaceae herbivores from the butterfly family Pieridae (*Pieris brassicae*, *P. macdunnoughii*, *P. napi* and *P. rapae*), one specialist herbivore, *Plutella xylostella* from Plutellidae and seven generalist herbivores from the Noctuidae (*Helicoverpa armigera*, *Helicoverpa zea*, *Mamestra configurata*, *Noctua pronuba*, *Trichoplusia ni*, *Spodoptera exigua* and *Phlogophora meticulosa*). BUSCO analysis of proteomes indicated that all the species had > 87.5% completeness with an average of 90.9% completeness, <5% average duplication, <1.5% average fragmentation and < 3.5% average missingness. The number of annotated protein coding genes ranged from 16733 in *P. napi* to 28532 proteins in *P. meticulosa* (figure. 1; mean: 20730 proteins). The number of annotated proteins was higher in moths (families Noctuidae and Plutellidae; median: 20557) compared to butterflies (family Pieridae: 17053 proteins). The number of functionally annotated proteins ranged from 12840 to 17850 (mean: 14650 annotations).

**Figure 1:**
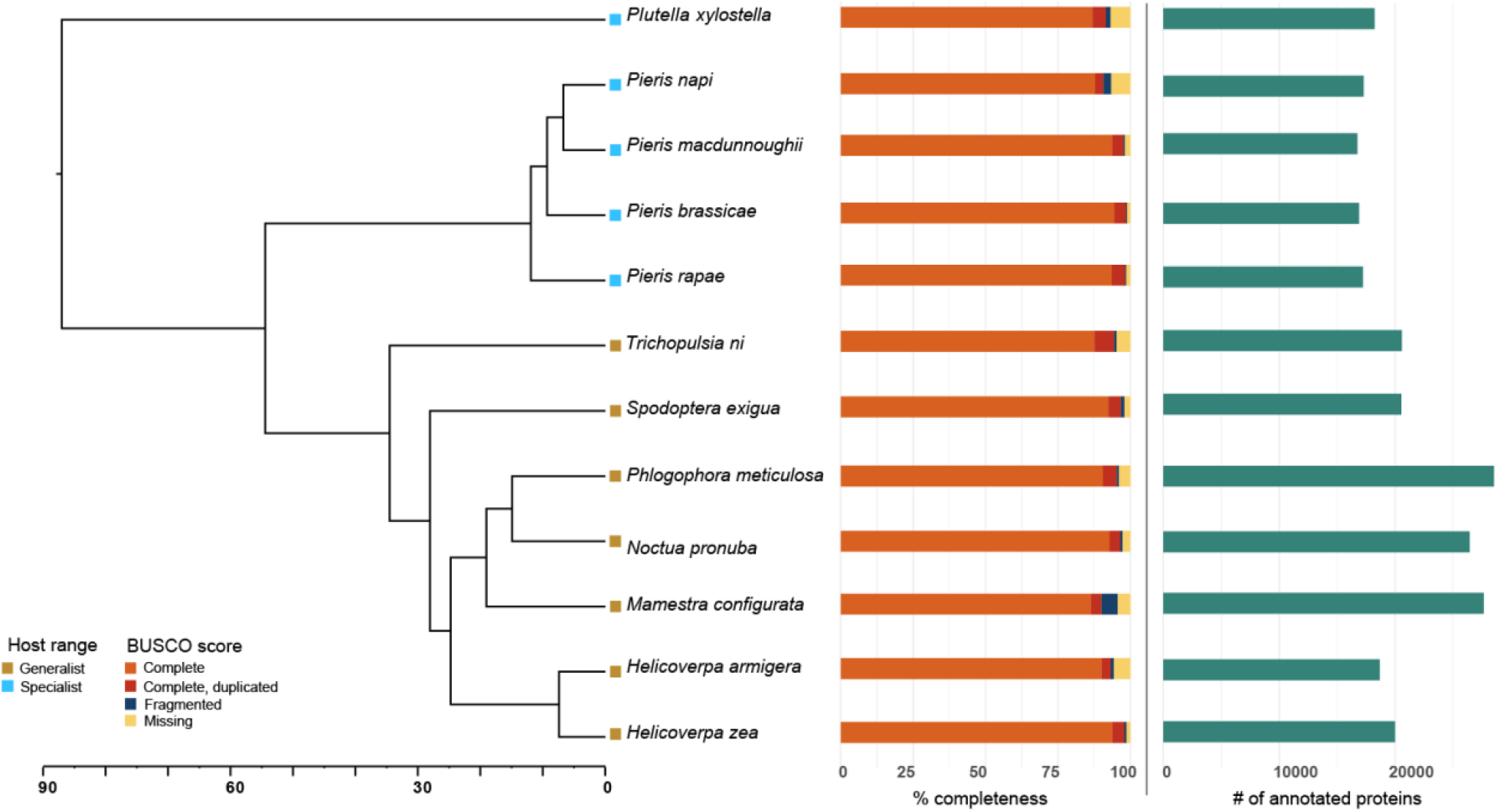
Maximum Likelihood tree based on single copy orthologs from 12 Lepidopteran species. Host range legend reflects herbivore specialist/generalist status. BUSCO score represents protein completeness.

OrthoFinder analysis identified 21785 orthogroups. The gene counts corresponding to these orthogroups were used as input for CAFE analysis. Our filtered OG count after removing high variance OGs among species and >100 genes in any single species included 20897 OGs in the “all genes” dataset. The “detoxification genes” dataset consisted of 690 OGs and the “sensory genes” dataset consisted of 69 OGs. The “single gene family” dataset consisted of the following number of OGs for each family: ABCs: 57, COesterases: 77, Insect cuticle proteins: 123, GSTs: 33, Gustatory genes: 6, Ionotropic genes: 14, Olfactory genes: 49, p450s: 101, Sulfotransferases: 28, Trypsins: 237 and finally, UGTs: 34.

Our species level phylogenetic tree was constructed based on 3575 complete single copy orthologs. Our phylogeny placed *Plutella xylostella* as the outgroup and Pieridae and Noctuidae as sister groups which is consistent with other published phylogenies (figure. 1).

### Gene family expansions and contractions

Our analysis suggested that across the entire phylogeny, *Mamestra configurata* had the highest gene family expansions for the “all genes” dataset (n=1873) and *Trichoplusia ni* had the highest gene family contractions (n=1269) (figure. 2). Among butterflies, *Pieris napi* had the highest gene family expansions (n=894) and gene family contractions (n=995) for the “all genes” dataset. Among the detoxification and sensory gene families, *M. configurata* had the highest gene family expansions for both families and the node leading to the *Pieris sps.* clade had the largest gene family contraction for detoxification genes and *P. napi* for the sensory gene families (figure. 2).

**Figure 2:**
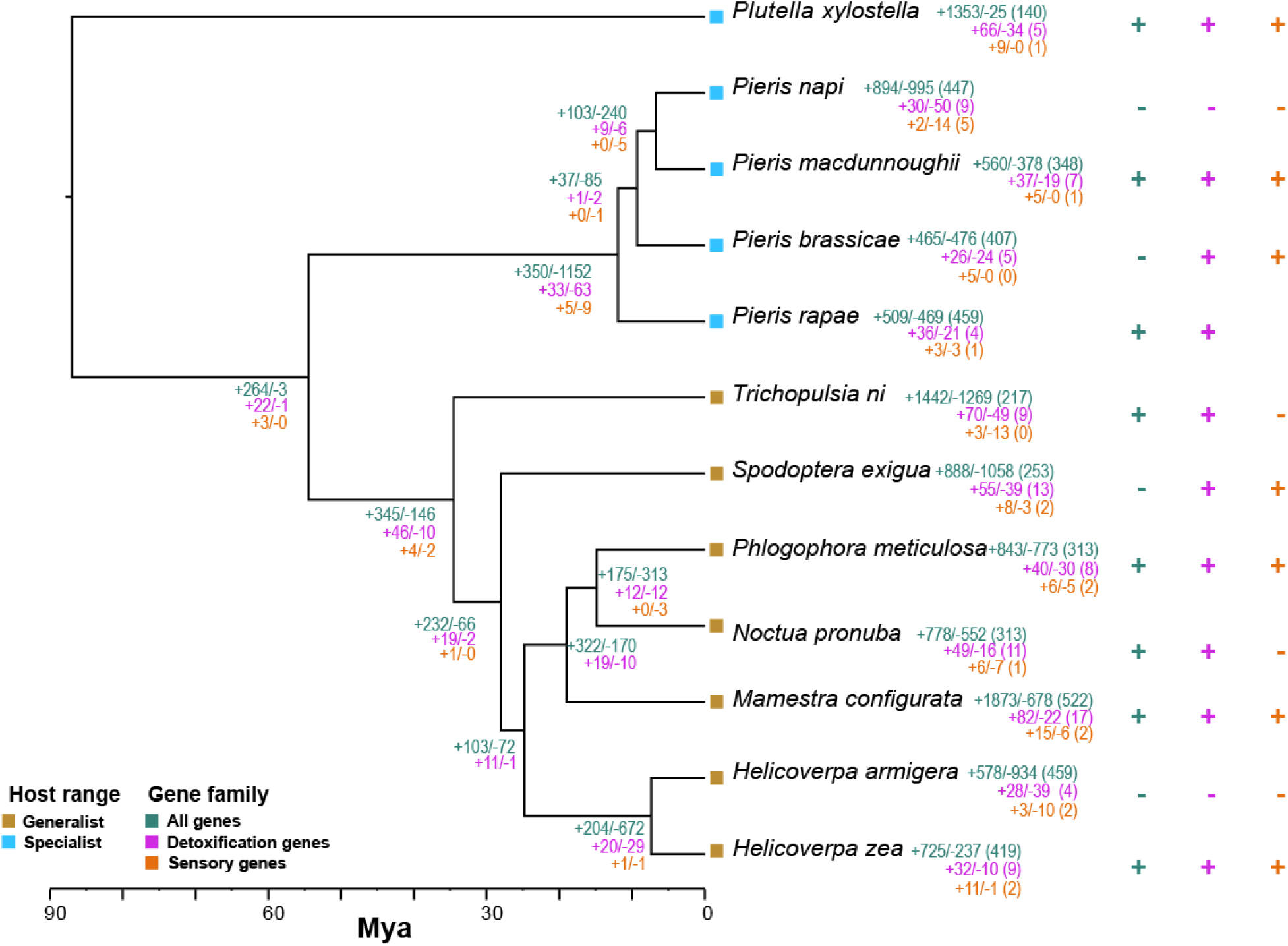
Maximum Likelihood tree showing gene family expansion (indicated as +) and contraction (indicated as -) for 12 species. Gene family color legend reflects expansion/contractions for the corresponding gene family.

### Gene family evolution

Our analysis of gene family evolution for the “all genes” dataset resulted in an overall change, λ, of 0.00274 (-logL: 114663; figure. 3). Separate λ estimations for “all genes” showed that gene family evolution in *P*. *xylostella* (λ= 0.00058) was lower than the baseline change, while both specialists (λ=0.0027) and generalists (λ=0.0036) had a higher rate of evolution than baseline, with generalists having the highest rate of gene evolution. For “detoxification genes”, generalist herbivores had higher rate of gene evolution (λ= 0.0071) compared to specialist herbivores (λ= 0.0062) and for “sensory genes”, specialist herbivores had higher gene evolution rate (λ= 0.0059) than generalist herbivores.

**Figure 3:**
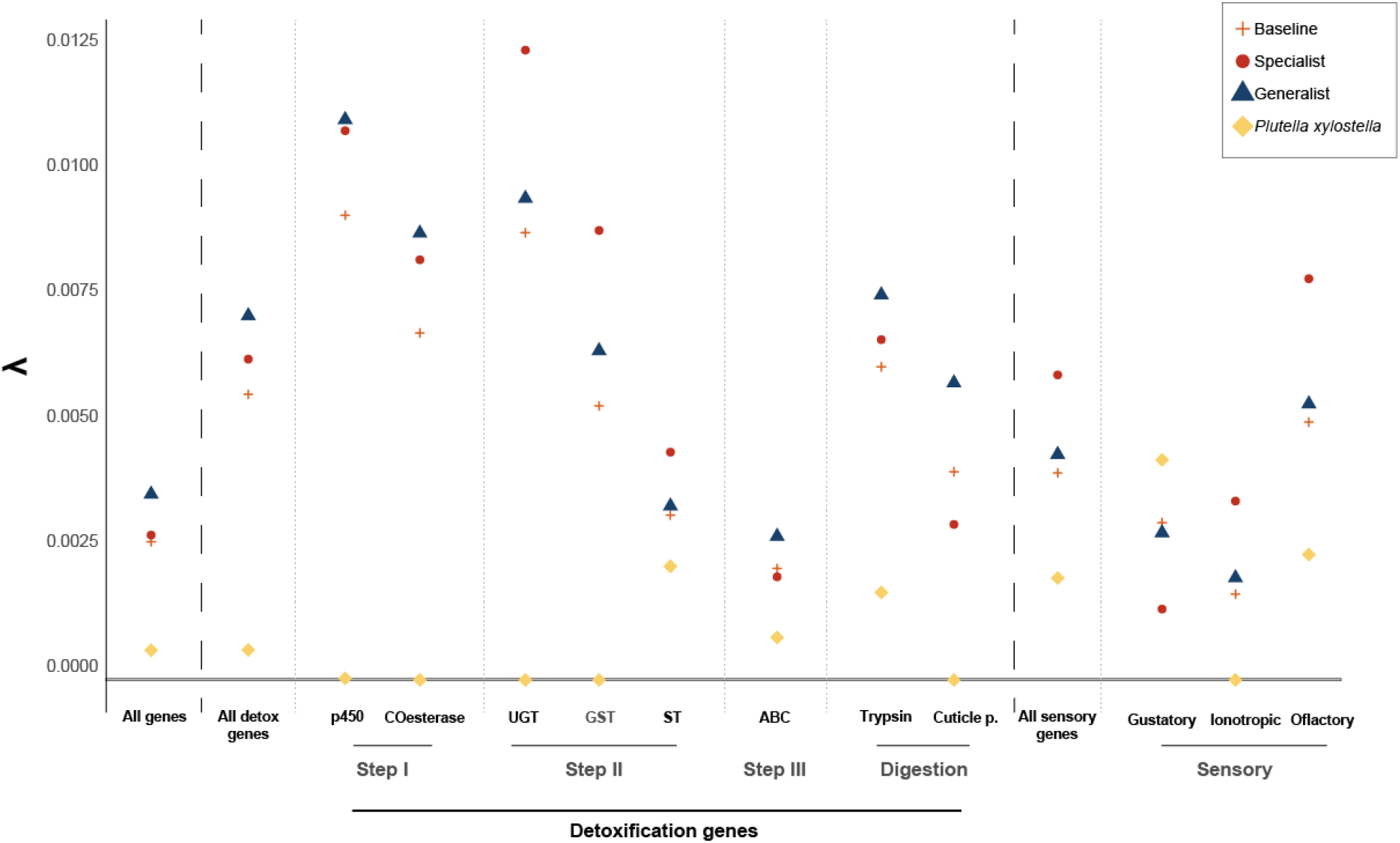
Gene family evolution rates calculated from CAFE. Rates reflect change/gene/Myr.

**Figure 4:**
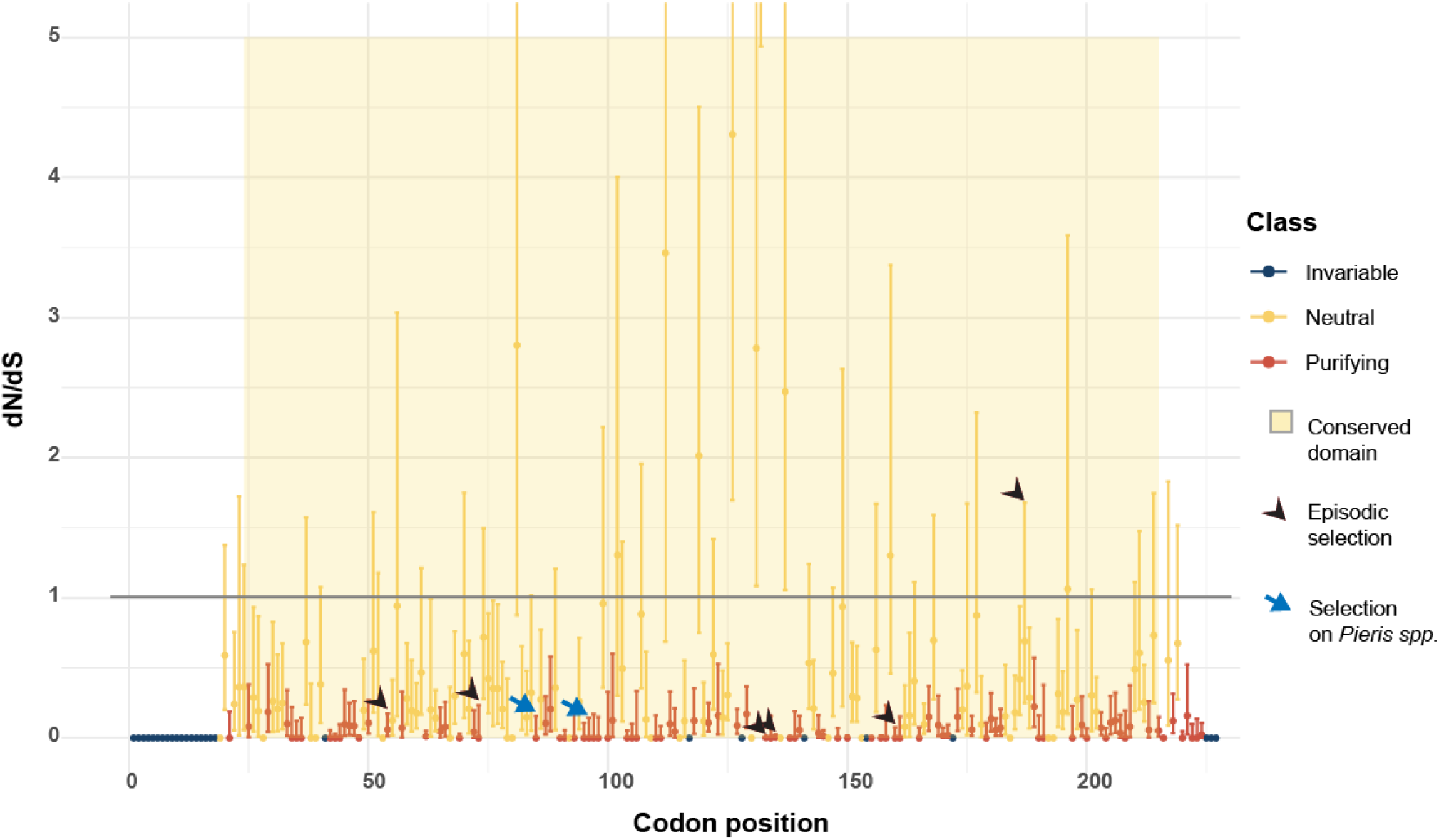
Molecular evolution of NSP gene. Episodic selection and differential selection highlighted in black and blue arrows respectively.

In our “single gene family” dataset, *P. xylostella* had the lowest rate of evolution across all detoxification enzymes. In genes that constitute the first step in detoxification of plant compounds, generalist herbivores had the highest rate of change in p450 genes (λ=0.011) and in COesterase gene family (λ=0.0087). In the genes that facilitate conversion of plant toxic compounds to hydrophilic compounds, specialist herbivores had the highest rate of evolution in UGT (λ=0.0123), GST (λ=0.0087) and ST (λ=0.004). Generalist herbivores had higher rate of evolution for ABC (λ=0.0028) (which is involved in excreting toxic compounds), for trypsin (λ=0.007) and cuticle protein (λ=0.0058) (both involved in increasing digestive efficiency).

Our analysis of “single gene family” olfactory gene families indicated that gustatory gene evolution was highest in *P. xylostella* (λ=0.0043) and lowest in specialist herbivores (λ=0.0013) (*Pieris spp.*) while ionotropic and olfactory gene families evolved at a faster rate in specialist herbivores compared generalist feeders or *P. xylostella* (figure. 3).

### Signatures of selection on NSP gene

We found evidence for pervasive purifying selection in the NSP gene (fig. 4.4). 41% of the codons were under purifying selection (95/227 codons). Our analysis of dN/dS showed that the NSP gene was characterized predominantly by synonymous substitutions. Almost all of the codons under purifying selection were present in the conserved domain of the NSP gene. However, we did not identify any sites that were under pervasive diversifying/positive selection. We used Contrast-FEL to test if the NSP gene in the *Pieris* genus had different substitution rates compared to the rest of the phylogeny. We identified 2 sites (blue arrow in fig. 4.4) that showed different substitution rates in the *Pieris spp.* clade compared to the rest of the phylogeny. Additionally, we identified six sites in the conserved domain that were under episodic positive/diversifying selection (black arrow in fig. 4.4).

## Discussion

We evaluated the role of detoxification and sensory gene family expansions in specialist and generalist herbivores feeding on Brassicaceae. We additionally examined selection on the nitrile specifier protein gene, a novel protein involved in glucosinolate detoxification in Lepidoptera. We found that generalist herbivores had the largest gene family changes in all gene families, detoxification related families as well as sensory gene families. Generalist herbivores also had faster evolution of all genes and detoxification gene families, while specialists had faster gene evolution in sensory gene families. Additionally, we found that the nitrile specifier protein gene is highly conserved among generalist and specialist lepidopterans. Although NSP is highly conserved, we identified signatures of episodic and positive selection on the gene.

### Gene family expansion/contraction in Lepidoptera

We examined gene family expansions/contractions in 12 species of Lepidoptera from 3 distinct families that feed on Brassicaceae. Although many other Lepidoptera and insects are known to feed on Brassicaceae, the 12 species included in our analysis are known to cause serious damage to Brassicaceae. In fact, species in the Noctuidae family constitute some of the most damaging pests in the world (Cho et al., 2008). Genera such as *Spodoptera*, *Trichoplusia* and *Helicoverpa* are some of the insects with the widest range of recorded hostplants (Robinson et al., 2010). All species in the Noctuidae family in our analysis are polyphagous and also feed on Brassicaceae, while Pieridae and Plutellidae family in our analysis consists of specialist herbivores on Brassicaceae.

We found that Noctuidae family and the species represented within showed the largest gene family expansions and contractions, followed by *Plutella xylostella* (figure. 2). *Pieris sps.* had the lowest gene family expansion. Our results, therefore, suggest that polyphagy is associated with greater gene family expansions and contractions. This is in line with other studies that have found that species with larger hostplant ranges have large gene families compared to those that fed on few or a single plant family (Breeschoten et al., 2022; Suzuki et al., 2018).

Polyphagous herbivores use the general detoxification processes (phase I -III) to detoxify plant compounds and thus, larger copy numbers of these genes will result in higher gene expression to process plant toxins efficiently (Breeschoten et al., 2022; Dermauw & Van Leeuwen, 2014; Ranson et al., 2002). Polyphagy is also associated with a dynamic gene family expansion/contraction, especially if the hostplant range of the herbivore is labile. For example, in the Nymphalidae, hostplant range oscillates from ancestral specialists that experience range extension giving rise to generalist feeders that adapt to local hostplant communities and in turn diversify and specialize (Hardy & Otto, 2014; Janz et al., 2006; Janz & Nylin, 2008; Nylin et al., 2014). This results in rapid changes in gene family expansions/contractions. If hostplant specialization is selected in the future, this can result in fewer gene family expansion/contractions. In fact, that is what we see in specialist Brassicaceae feeders. *Pieris spp.* and *P. xylostella* on the other hand are specialist herbivores that possess fine-tuned detoxification processes to counteract glucosinolate toxicity. Therefore, generalized detoxification genes families are relatively stable and under strong selection.

### Gene evolution in Lepidoptera

Our CAFE analysis estimated an overall change of 0.00274 (gains/losses/Myr) for all genes. Our estimate of λ is consistent with what is seen in other insects such as *Drosophila*, λ= 0.0012; (Hahn et al., 2007), Lepidoptera λ= 0.0023; (Breeschoten et al., 2022), and *Anopheles* λ= 0.0031 (Neafsey et al., 2015). When estimating separate change for specialists, generalists, and *P. xylostella*, we found a higher rate of change for generalists, which fits with the largest gene family expansion/contraction in generalist species (Breeschoten et al., 2022; Suzuki et al., 2018). Higher gene family evolution rates could be due to underlying gene duplication, neo-functionalization and/or genome rearrangements, all of which are implicated in polyphagous feeding (Murad et al., 2021; Seppey et al., 2019).

Our estimation of gene evolution for detoxification gene families showed that generalists had the highest rate of change. When we examined individual detoxification gene families, generalists had higher gene evolution for phase I, phase III and digestion related genes (cuticle and trypsin). p450 genes have been extensively studied for their role in hostplant use in Lepidoptera. p450 genes can metabolize a wide range of plant compounds including xanthoxin (Mao et al., 2006), furanocumarins (Lindroth, 1989), alkaloids (Wang et al., 2018) and even insecticide resistance (Pan et al., 2018). Since polyphagous herbivores feed on a wide range of chemicals, faster evolution in genes of phase I detoxification can help them adapt to new hosts and overcome insecticide resistance. Gene families in phase II had a higher evolutionary rate in specialist herbivores compared to generalists. Phase II detoxification genes are specialized genes that add specific compounds which increase solubility and excretion efficiency (Nallu et al., 2018; You et al., 2015). Specialist herbivores have alternate pathways to detoxify toxic compounds, for example, the use of NSP to form nitriles instead of isothiocyanates in specialist Brassicaceae feeders (Edger et al., 2015; Okamura, Sato, Tsuzuki, Sawada, et al., 2019; Ratzka et al., 2002). These modified products are processed by phase II genes and specialist herbivores have evolved modified phase II enzymes to efficiently process these by products. In fact, GSTs are upregulated in the larvae of *Pieris spp.* and *Plutella xylostella* when feeding on Brassicaceae (You et al., 2015). UDPGTs are also enriched in the transcriptome of *Danus plexippus* larvae which feed on plants from the family Apocynaceae which contain cardiac glycosides (Ranz et al., 2021). Generalist herbivores had higher evolution in phase III as well trypsin and insect cuticle proteins, all of which are positively correlated with polyphagy in many insect species including Lepidoptera (Kelkenberg et al., 2015; Muhlia-Almazán et al., 2008; Rawlings & Barrett, 1995).

Our estimation of gene evolution in sensory receptors highlighted that overall, specialist herbivores had faster evolution compared to generalists. Our single gene family analysis, however showed faster evolution of gustatory receptors in *P. xylostella* compared to generalists or specialists. This is in agreement with what has been published for lepidopteran gustatory receptors. *Plutella xylostella* is an outlier and possess greater diversity of gustatory receptors and is known to have higher gene evolution compared to other lepidopteran species (Engsontia et al., 2014). The gene evolution of gustatory gene families is positively correlated with host plant range. For example, in a transcriptomic analysis of gustatory gene evolution in Nymphalidae, the lineage leading to *Vanessa cardui*, a generalist herbivore known to feed on over 400 hostplants had the highest gene evolution rate compared to other specialists (Suzuki et al., 2018). Our analysis of ionotropic and olfactory gene evolution suggested higher rate of evolution in specialist herbivores compared to generalist herbivores. Specialist herbivores feeding on Brassicaceae have fine-tuned olfactory receptors to identify glucosinolates that have evolved through gene duplication and neofunctionalization and therefore expected to have higher evolution compared to those in generalist herbivores (Matsunaga et al., 2022).

### Selection on NSP gene

Our analysis of selection on the NSP gene in Brassicaceae-feeding Lepidopteran insects revealed strong purifying selection. Our dN/dS estimates indicated a lack of non-synonymous substitutions, suggesting the absence of pervasive diversifying selection on NSP. Our contrast-FEL analysis showed that the lineage leading to *Pieris spp.* had 2 sites in the conserved domain that had different substitution rates compared to the other species. This could indicate selection specific to *Pieris spp.* on the NSP gene. Our results are in agreement with what has been shown for positive selection on NSP gene in a subgroup of *Pieris spp*. (Okamura, Sato, Tsuzuki, Murakami, et al., 2019). The neofunctionalization and cooption of the insect allergen protein in *Pieris* spp. to the extant NSP gene is a key innovation that allowed *Pieris spp.* to colonize Brassicaceae and diversify on them. The two codons that we identified could represent the differential selection pressures experienced by *Pieris spp* compared to the rest of the phylogeny. Finally, our test for episodic selection identified 6 sites that were subjected to diversifying selection. Although our analysis does not shed light on the functional role of the sites under selection, it does highlight the likely genetic targets that are crucial for adaptation and feeding on Brassicaceae. These sites could serve as the targets for manipulative CRISPR experiments to understand the mechanisms through which specialist herbivores detoxify glucosinolates in Brassicaceae.

### Genome quality and Lepidopteran phylogenetic framework

Our interpretations of gene family evolution and selection on the NSP gene is critically dependent on the accuracy of genome annotations as well as species phylogeny. The availability of genomes for non-model organisms has increased considerably with the advent of affordable sequencing and development of assembly and annotation pipelines. However, the quality of assembly and annotation varies considerably due to differences in sequencing methods as well as differences in analytical methods among various research groups (Ellis et al., 2021). To control for variation among different genomes, we employed several strict quality checks and additional analysis: 1) We checked the quality of the genome using BUSCO and used only those genomes that had a BUSCO complete, single copy score >80%. 2.) We used BRAKER 2 for gene prediction and EggNOG for functional annotation instead of using the default proteome files available with the genomes. The annotated proteomes were assessed for quality using BUSCO and only those that had a BUSCO complete single copy score >85% were retained. This ensured that all proteomes were of comparable quality for downstream analysis. 3) We retained only the longest isoform per gene and removed isoform duplications that could bias gene family evolution estimates. 4) We assigned a separate lambda for Plutella xylostella since it was the outgroup in our analysis as well as a specialist Lepidoptera belonging to a different family from the other species. 5) We calculated the error in CAFE analysis and applied that to our analysis to account for genome annotation and assembly errors.

Our phylogenetic reconstruction using single copy orthologs placed *P. xylostella* as the outgroup with Noctuidae and Pieridae as sister genera which is consistent with published reports (Espeland et al., 2018). We used 87 MYA as the root calibration, the age of diversification of *P. xylostella*. This placed the age of node split of *P. napi* and *P. macdunnoughii* about 4-6 MYA, which is consistent with what the Holarctic expansion and diversification of *P. napi* (Geiger & Shapiro, 1992).

## Conclusion

We used whole genome comparative analysis to examine the role of detoxification gene families, sensory gene families and the specialized NSP gene on Lepidopteran herbivory on Brassicaceae. We found positive correlations with general detoxification gene expansions and polyphagy and identified specific steps in the detoxification pathway where gene family expansions were correlated with host specialization. We additionally identified signatures of pervasive purifying selection on the NSP gene as well as identified sites that were subjected to different selection pressures in the lineage leading to *Pieris* spp. Our results add to the growing body of work on understanding gene family evolution and its role in hostplant range and specialization in insects.

## Acknowledgements

We thank Ward B Watt, Joseph M Quattro, Brian Hollis, and Peter Andolfatto for proving comments on earlier versions of the manuscript. We thank Muktai Kuwalekar, and Hannah Dort for advice on gene family evolution analysis. This work was supported by the University of South Carolina Arts & Sciences to CLB.

## Notes

### Competing Interest Statement

The authors have declared no competing interest.

## References

Awmack, C. S., & Leather, S. R. (2002). Host Plant Quality and Fecundity in Herbivorous Insects. Annual Review of Entomology, 47(1), 817–844. https://doi.org/10.1146/annurev.ento.47.091201.145300

Berenbaum, M. (1983). Coumarins and Caterpillars: A Case for Coevolution. Evolution, 37(1), 163–179. https://doi.org/10.2307/2408184

Berenbaum, M. R., Zangerl, A. R., & Nitao, J. K. (1986). Constraints on Chemical Coevolution: Wild Parsnips and the Parsnip Webworm. Evolution, 40(6), 1215–1228. https://doi.org/10.1111/j.1558-5646.1986.tb05746.x

Bernays, E., & Graham, M. (1988). On the Evolution of Host Specificity in Phytophagous Arthropods. Ecology, 69(4), 886–892. https://doi.org/10.2307/1941237

Brand, P., Robertson, H. M., Lin, W., Pothula, R., Klingeman, W. E., Jurat-Fuentes, J. L., & Johnson, B. R. (2018). The origin of the odorant receptor gene family in insects. ELife, 7, e38340. https://doi.org/10.7554/eLife.38340

Breeschoten, T., van der Linden, C. F. H., Ros, V. I. D., Schranz, M. E., & Simon, S. (2022). Expanding the Menu: Are Polyphagy and Gene Family Expansions Linked across Lepidoptera? Genome Biology and Evolution, 14(1). https://doi.org/10.1093/gbe/evab283

Brůna, T., Hoff, K. J., Lomsadze, A., Stanke, M., & Borodovsky, M. (2021). BRAKER2: Automatic eukaryotic genome annotation with GeneMark-EP+ and AUGUSTUS supported by a protein database. NAR Genomics and Bioinformatics, 3(1), lqaa108. https://doi.org/10.1093/nargab/lqaa108

Calla, B., Noble, K., Johnson, R. M., Walden, K. K. O., Schuler, M. A., Robertson, H. M., & Berenbaum, M. R. (2017). Cytochrome P450 diversification and hostplant utilization patterns in specialist and generalist moths: Birth, death and adaptation. Molecular Ecology, 26(21), 6021–6035. https://doi.org/10.1111/mec.14348

Cantalapiedra, C. P., Hernández-Plaza, A., Letunic, I., Bork, P., & Huerta-Cepas, J. (2021). eggNOG-mapper v2: Functional Annotation, Orthology Assignments, and Domain Prediction at the Metagenomic Scale. Molecular Biology and Evolution, 38(12), 5825– 5829. https://doi.org/10.1093/molbev/msab293

Cermak, R. (2008). Effect of dietary flavonoids on pathways involved in drug metabolism. Expert Opinion on Drug Metabolism & Toxicology, 4(1), 17–35. https://doi.org/10.1517/17425255.4.1.17

Challi, R. J., Kumar, S., Dasmahapatra, K. K., Jiggins, C. D., & Blaxter, M. (2016). Lepbase: The Lepidopteran genome database. BioRxiv. https://doi.org/10.1101/056994

Chew, F. S. (1977). Coevolution of Pierid Butterflies and Their Cruciferous Foodplants. II. The Distribution of Eggs on Potential Foodplants. Evolution, 31(3), 568–579. https://doi.org/10.2307/2407522

Chew, F. S. (1980). Foodplant preferences of Pieris caterpillars (Lepidoptera). Oecologia, 46(3), 347–353. https://doi.org/10.1007/BF00346263

Cho, S., Mitchell, A., Mitter, C., Regier, J., Matthews, M., & Robertson, R. (2008). Molecular phylogenetics of heliothine moths (Lepidoptera: Noctuidae: Heliothinae), with comments on the evolution of host range and pest status. Systematic Entomology, 33(4), 581–594. https://doi.org/10.1111/j.1365-3113.2008.00427.x

Dainat, J., Hereñú, D., & Pucholt, P. (2020). AGAT: Another Gff Analysis Toolkit to handle annotations in any GTF. GFF Format. Zenodo.

Dermauw, W., & Van Leeuwen, T. (2014). The ABC gene family in arthropods: Comparative genomics and role in insecticide transport and resistance. Insect Biochemistry and Molecular Biology, 45, 89–110. https://doi.org/10.1016/j.ibmb.2013.11.001

Dermauw, W., Van Leeuwen, T., & Feyereisen, R. (2020). Diversity and evolution of the P450 family in arthropods. Insect Biochemistry and Molecular Biology, 127, 103490. https://doi.org/10.1016/j.ibmb.2020.103490

Edger, P. P., Heidel-Fischer, H. M., Bekaert, M., Rota, J., Glöckner, G., Platts, A. E., Heckel, D. G., Der, J. P., Wafula, E. K., Tang, M., Hofberger, J. A., Smithson, A., Hall, J. C., Blanchette, M., Bureau, T. E., Wright, S. I., dePamphilis, C. W., Eric Schranz, M., Barker, M. S., … Wheat, C. W. (2015). The butterfly plant arms-race escalated by gene and genome duplications. Proceedings of the National Academy of Sciences, 112(27), 8362–8366. https://doi.org/10.1073/pnas.1503926112

Ehrlich, P. R., & Raven, P. H. (1964). Butterflies and Plants: A Study in Coevolution. Evolution, 18(4), 586–608.

Ellis, E. A., Storer, C. G., & Kawahara, A. Y. (2021). De novo genome assemblies of butterflies. GigaScience, 10(6), 1–8. https://doi.org/10.1093/gigascience/giab041

Emms, D. M., & Kelly, S. (2019). OrthoFinder: Phylogenetic orthology inference for comparative genomics. Genome Biology, 20(1), 238. https://doi.org/10.1186/s13059-019-1832-y

Engsontia, P., Sangket, U., Chotigeat, W., & Satasook, C. (2014). Molecular evolution of the odorant and gustatory receptor genes in lepidopteran insects: Implications for their adaptation and speciation. Journal of Molecular Evolution, 79(1–2), 21–39. https://doi.org/10.1007/s00239-014-9633-0

Espeland, M., Breinholt, J., Willmott, K. R., Warren, A. D., Vila, R., Toussaint, E. F. A., Maunsell, S. C., Aduse-Poku, K., Talavera, G., Eastwood, R., Jarzyna, M. A., Guralnick, R., Lohman, D. J., Pierce, N. E., & Kawahara, A. Y. (2018). A comprehensive and dated phylogenomic analysis of butterflies. Current Biology, 28(5), 770--778.e5. https://doi.org/10.1016/j.cub.2018.01.061

Feyereisen, R. (1999). Insect P450 Enzymes. Annual Review of Entomology, 44(1), 507–533. https://doi.org/10.1146/annurev.ento.44.1.507

Futuyma, D. J., & Agrawal, A. A. (2009). Macroevolution and the biological diversity of plants and herbivores. Proceedings of the National Academy of Sciences, 106(43), 18054– 18061. https://doi.org/10.1073/pnas.0904106106

Geiger, H., & Shapiro, A. M. (1992). Genetics, systematics and evolution of holarctic *Pieris napi* species group populations (Lepidoptera, Pieridae). Journal of Zoological Systematics and Evolutionary Research, 30(2), 100–122. https://doi.org/10.1111/j.1439-0469.1992.tb00161.x

Girgis, H. Z. (2015). Red: An intelligent, rapid, accurate tool for detecting repeats de-novo on the genomic scale. BMC Bioinformatics, 16(1), 227. https://doi.org/10.1186/s12859-015-0654-5

Gripenberg, S., Mayhew, P. J., Parnell, M., & Roslin, T. (2010). A meta-analysis of preference-performance relationships in phytophagous insects. Ecology Letters, 13(3), 383–393. https://doi.org/10.1111/j.1461-0248.2009.01433.x

Hahn, M. W., Han, M. V., & Han, S. G. (2007). Gene family evolution across 12 Drosophila genomes. PLoS Genetics, 3(11), 2135–2146. https://doi.org/10.1371/JOURNAL.PGEN.0030197

Halkier, B. A., & Gershenzon, J. (2006). Biology and Biochemistry of Glucosinolates. Annual Review of Plant Biology, 57(1), 303–333. https://doi.org/10.1146/annurev.arplant.57.032905.105228

Halon, E., Eakteiman, G., Moshitzky, P., Elbaz, M., Alon, M., Pavlidi, N., Vontas, J., & Morin, S. (2015). Only a minority of broad-range detoxification genes respond to a variety of phytotoxins in generalist *Bemisia tabaci* species. Scientific Reports, 5(1), Article 1. https://doi.org/10.1038/srep17975

Hardy, N. B., & Otto, S. P. (2014). Specialization and generalization in the diversification of phytophagous insects: Tests of the musical chairs and oscillation hypotheses. Proceedings of the Royal Society B: Biological Sciences, 281(1795), 20132960. https://doi.org/10.1098/rspb.2013.2960

Heidel-Fischer, H. M., & Vogel, H. (2015). Molecular mechanisms of insect adaptation to plant secondary compounds. Current Opinion in Insect Science, 8, 8–14. https://doi.org/10.1016/j.cois.2015.02.004

Jaenike, J. (1990). Host Specialization in Phytophagous Insects. Annual Review of Ecology and Systematics, 21, 243–273.

Janz, N., & Nylin, S. (2008). The oscillation hypothesis of host-plant range and speciation. Specialization, Speciation, and Radiation: The Evolutionary Biology of Herbivorous Insects. University of California Press Berkeley.

Janz, N., Nylin, S., & Wahlberg, N. (2006). Diversity begets diversity: Host expansions and the diversification of plant-feeding insects. BMC Evolutionary Biology, 6(1), 4. https://doi.org/10.1186/1471-2148-6-4

Jones, P., Binns, D., Chang, H.-Y., Fraser, M., Li, W., McAnulla, C., McWilliam, H., Maslen, J., Mitchell, A., Nuka, G., Pesseat, S., Quinn, A. F., Sangrador-Vegas, A., Scheremetjew, M., Yong, S.-Y., Lopez, R., & Hunter, S. (2014). InterProScan 5: Genome-scale protein function classification. Bioinformatics, 30(9), 1236–1240. https://doi.org/10.1093/bioinformatics/btu031

Joshi, A., & Thompson, J. N. (1995). Trade-offs and the evolution of host specialization. Evolutionary Ecology, 9(1), 82–92. https://doi.org/10.1007/BF01237699

Kant, M. R., Jonckheere, W., Knegt, B., Lemos, F., Liu, J., Schimmel, B. C. J., Villarroel, C. A., Ataide, L. M. S., Dermauw, W., Glas, J. J., Egas, M., Janssen, A., Van Leeuwen, T., Schuurink, R. C., Sabelis, M. W., & Alba, J. M. (2015). Mechanisms and ecological consequences of plant defence induction and suppression in herbivore communities. Annals of Botany, 115(7), 1015–1051. https://doi.org/10.1093/aob/mcv054

Katoh, K., & Standley, D. M. (2013). MAFFT Multiple Sequence Alignment Software Version 7: Improvements in Performance and Usability. Molecular Biology and Evolution, 30(4), 772–780. https://doi.org/10.1093/molbev/mst010

Kelkenberg, M., Odman-Naresh, J., Muthukrishnan, S., & Merzendorfer, H. (2015). Chitin is a necessary component to maintain the barrier function of the peritrophic matrix in the insect midgut. Insect Biochemistry and Molecular Biology, 56, 21–28. https://doi.org/10.1016/j.ibmb.2014.11.005

Kriventseva, E. V., Kuznetsov, D., Tegenfeldt, F., Manni, M., Dias, R., Simão, F. A., & Zdobnov, E. M. (2019). OrthoDB v10: Sampling the diversity of animal, plant, fungal, protist, bacterial and viral genomes for evolutionary and functional annotations of orthologs. Nucleic Acids Research, 47(D1), D807–D811. https://doi.org/10.1093/nar/gky1053

Kuchernig, J. C., Burow, M., & Wittstock, U. (2012). Evolution of specifier proteins in glucosinolate-containing plants. BMC Evolutionary Biology, 12(127). http://www.biomedcentral.com/1471-2148/12/127

Lindroth, R. L. (1989). Host plant alteration of detoxication activity in *Papilio glaucus glaucus*. Entomologia Experimentalis et Applicata, 50(1), 29–35. https://doi.org/10.1111/j.1570-7458.1989.tb02310.x

Mao, W., Berhow, M. A., Zangerl, A. R., Mcgovern, J., & Berenbaum, M. R. (2006). Cytochrome P450-Mediated Metabolism of Xanthotoxin by *Papilio multicaudatus*. Journal of Chemical Ecology, 32(3), 523–536. https://doi.org/10.1007/s10886-005-9018-3

Matsunaga, T., Reisenman, C. E., Goldman-Huertas, B., Brand, P., Miao, K., Suzuki, H. C., Verster, K. I., Ramírez, S. R., & Whiteman, N. K. (2022). Evolution of Olfactory Receptors Tuned to Mustard Oils in Herbivorous Drosophilidae. Molecular Biology and Evolution, 39(2), msab362. https://doi.org/10.1093/molbev/msab362

Mendes, F. K., Vanderpool, D., Fulton, B., & Hahn, M. W. (2020). CAFE 5 models variation in evolutionary rates among gene families. Bioinformatics, 36(22–23), 5516–5518. https://doi.org/10.1093/bioinformatics/btaa1022

Missbach, C., Dweck, H. K., Vogel, H., Vilcinskas, A., Stensmyr, M. C., Hansson, B. S., & Grosse-Wilde, E. (2014). Evolution of insect olfactory receptors. ELife, 3, e02115. https://doi.org/10.7554/eLife.02115

Mistry, J., Chuguransky, S., Williams, L., Qureshi, M., Salazar, G. A., Sonnhammer, E. L. L., Tosatto, S. C. E., Paladin, L., Raj, S., Richardson, L. J., Finn, R. D., & Bateman, A. (2021). Pfam: The protein families database in 2021. Nucleic Acids Research, 49(D1), D412–D419. https://doi.org/10.1093/nar/gkaa913

Muhlia-Almazán, A., Sánchez-Paz, A., & García-Carreño, F. L. (2008). Invertebrate trypsins: A review. Journal of Comparative Physiology B, 178(6), 655–672. https://doi.org/10.1007/s00360-008-0263-y

Murad, N. F., Silva-Brandão, K. L., & Brandão, M. M. (2021). Mechanisms behind polyphagia in a pest insect: Responses of *Spodoptera frugiperda* (J.E. Smith) strains to preferential and alternative larval host plants assessed with gene regulatory networks. Biochimica et Biophysica Acta (BBA) – Gene Regulatory Mechanisms, 1864(3), 194687. https://doi.org/10.1016/j.bbagrm.2021.194687

Nallu, S., Hill, J. A., Don, K., Sahagun, C., Zhang, W., Meslin, C., Snell-Rood, E., Clark, N. L., Morehouse, N. I., Bergelson, J., Wheat, C. W., & Kronforst, M. R. (2018). The molecular genetic basis of herbivory between butterflies and their host plants. Nature Ecology & Evolution, 2(9), Article 9. https://doi.org/10.1038/s41559-018-0629-9

Neafsey, D. E., Waterhouse, R. M., Abai, M. R., Aganezov, S. S., Alekseyev, M. A., Allen, J. E., Amon, J., Arcà, B., Arensburger, P., Artemov, G., Assour, L. A., Basseri, H., Berlin, A., Birren, B. W., Blandin, S. A., Brockman, A. I., Burkot, T. R., Burt, A., Chan, C. S., … Besansky, N. J. (2015). Highly evolvable malaria vectors: The genomes of 16 Anopheles mosquitoes. Science, 347(6217), 1258522. https://doi.org/10.1126/science.1258522

Nei, M., Niimura, Y., & Nozawa, M. (2008). The evolution of animal chemosensory receptor gene repertoires: Roles of chance and necessity. Nature Reviews Genetics, 9(12), Article 12. https://doi.org/10.1038/nrg2480

Nguyen, L.-T., Schmidt, H. A., von Haeseler, A., & Minh, B. Q. (2015). IQ-TREE: A Fast and Effective Stochastic Algorithm for Estimating Maximum-Likelihood Phylogenies. Molecular Biology and Evolution, 32(1), 268–274. https://doi.org/10.1093/molbev/msu300

Nylin, S., Slove, J., & Janz, N. (2014). Host Plant Utilization, Host Range Oscillations And Diversification In Nymphalid Butterflies: A Phylogenetic Investigation. Evolution, 68(1), 105–124. https://doi.org/10.1111/evo.12227

Okamura, Y., Sato, A., Tsuzuki, N., Murakami, M., Heidel-Fischer, H., & Vogel, H. (2019). Molecular signatures of selection associated with host plant differences in *Pieris* butterflies. Molecular Ecology, 28(22), 4958–4970. https://doi.org/10.1111/mec.15268

Okamura, Y., Sato, A., Tsuzuki, N., Sawada, Y., Hirai, M. Y., Heidel-Fischer, H., Reichelt, M., Murakami, M., & Vogel, H. (2019). Differential regulation of host plant adaptive genes in *Pieris* butterflies exposed to a range of glucosinolate profiles in their host plants. Scientific Reports, 9(1), Article 1. https://doi.org/10.1038/s41598-019-43703-8

Ozaki, K., Ryuda, M., Yamada, A., Utoguchi, A., Ishimoto, H., Calas, D., Marion-Poll, F., Tanimura, T., & Yoshikawa, H. (2011). A gustatory receptor involved in host plant recognition for oviposition of a swallowtail butterfly. Nature Communications, 2(1), Article 1. https://doi.org/10.1038/ncomms1548

Pan, Y., Chai, P., Zheng, C., Xu, H., Wu, Y., Gao, X., Xi, J., & Shang, Q. (2018). Contribution of cytochrome P450 monooxygenase CYP380C6 to spirotetramat resistance in *Aphis gossypii* Glover. Pesticide Biochemistry and Physiology, 148, 182–189. https://doi.org/10.1016/j.pestbp.2018.04.015

Pomés, A., Melén, E., Vailes, L. D., Retief, J. D., Arruda, L. K., & Chapman, M. D. (1998). Novel Allergen Structures with Tandem Amino Acid Repeats Derived from German and American Cockroach. Journal of Biological Chemistry, 273(46), 30801–30807. https://doi.org/10.1074/jbc.273.46.30801

Pond, S. L. K., Frost, S. D. W., & Muse, S. V. (2005). HyPhy: Hypothesis testing using phylogenies. *Bioinformatics (Oxford*, England), 21(5), 676–679. https://doi.org/10.1093/bioinformatics/bti079

Rane, R. V., Ghodke, A. B., Hoffmann, A. A., Edwards, O. R., Walsh, T. K., & Oakeshott, J. G. (2019). Detoxifying enzyme complements and host use phenotypes in 160 insect species. Current Opinion in Insect Science, 31, 131–138. https://doi.org/10.1016/j.cois.2018.12.008

Ranson, H., Claudianos, C., Ortelli, F., Abgrall, C., Hemingway, J., Sharakhova, M. V., Unger, M. F., Collins, F. H., & Feyereisen, R. (2002). Evolution of supergene families associated with insecticide resistance. Science, 298(5591), 179–181. https://doi.org/10.1126/science.1076781

Ranz, J. M., González, P. M., Clifton, B. D., Nazario-Yepiz, N. O., Hernández-Cervantes, P. L., Palma-Martínez, M. J., Valdivia, D. I., Jiménez-Kaufman, A., Lu, M. M., Markow, T. A., & Abreu-Goodger, C. (2021). A de novo transcriptional atlas in *Danaus plexippus* reveals variability in dosage compensation across tissues. Communications Biology, 4(1), Article 1. https://doi.org/10.1038/s42003-021-02335-3

Ratzka, A., Vogel, H., Kliebenstein, D. J., Mitchell-Olds, T., & Kroymann, J. (2002). Disarming the mustard oil bomb. Proceedings of the National Academy of Sciences, 99(17), 11223– 11228. https://doi.org/10.1073/pnas.172112899

Rawlings, N., & Barrett, A. (1995). Evolutionary families of metallopeptidases. 574 *Methods Enzymol*.

Robinson, G. S., Ackery, P. R., Kitching, I. J., Beccaloni, G. W., & Hernández, L. M. (2010). HOSTS – A Database of the World’s Lepidopteran Hostplants. Natural History Museum. https://www.nhm.ac.uk/our-science/data/hostplants/index.html

Rytz, R., Croset, V., & Benton, R. (2013). Ionotropic Receptors (IRs): Chemosensory ionotropic glutamate receptors in Drosophila and beyond. Insect Biochemistry and Molecular Biology, 43(9), 888–897. https://doi.org/10.1016/j.ibmb.2013.02.007

Ryuda, M., Calas-List, D., Yamada, A., Marion-Poll, F., Yoshikawa, H., Tanimura, T., & Ozaki, K. (2013). Gustatory Sensing Mechanism Coding for Multiple Oviposition Stimulants in the Swallowtail Butterfly, *Papilio xuthus*. Journal of Neuroscience, 33(3), 914–924. https://doi.org/10.1523/JNEUROSCI.1405-12.2013

Sánchez-Gracia, A., Vieira, F. G., & Rozas, J. (2009). Molecular evolution of the major chemosensory gene families in insects. Heredity, 103(3), 208–216. https://doi.org/10.1038/hdy.2009.55

Sayers, E. W., Beck, J., Bolton, E. E., Bourexis, D., Brister, J. R., Canese, K., Comeau, D. C., Funk, K., Kim, S., Klimke, W., Marchler-Bauer, A., Landrum, M., Lathrop, S., Lu, Z., Madden, T. L., O’Leary, N., Phan, L., Rangwala, S. H., Schneider, V. A., … Sherry, S. T. (2021). Database resources of the National Center for Biotechnology Information. Nucleic Acids Research, 49(D1), D10–D17. https://doi.org/10.1093/nar/gkaa892

Seppey, M., Ioannidis, P., Emerson, B. C., Pitteloud, C., Robinson-Rechavi, M., Roux, J., Escalona, H. E., McKenna, D. D., Misof, B., Shin, S., Zhou, X., Waterhouse, R. M., & Alvarez, N. (2019). Genomic signatures accompanying the dietary shift to phytophagy in polyphagan beetles. Genome Biology, 20(1), 98. https://doi.org/10.1186/s13059-019-1704-5

Simão, F. A., Waterhouse, R. M., Ioannidis, P., Kriventseva, E. V., & Zdobnov, E. M. (2015). BUSCO: Assessing genome assembly and annotation completeness with single-copy orthologs. Bioinformatics, 31(19), 3210–3212. https://doi.org/10.1093/bioinformatics/btv351

Suyama, M., Torrents, D., & Bork, P. (2006). PAL2NAL: Robust conversion of protein sequence alignments into the corresponding codon alignments. Nucleic Acids Research, 34(suppl_2), W609–W612. https://doi.org/10.1093/nar/gkl315

Suzuki, H. C., Ozaki, K., Makino, T., Uchiyama, H., Yajima, S., & Kawata, M. (2018). Evolution of Gustatory Receptor Gene Family Provides Insights into Adaptation to Diverse Host Plants in Nymphalid Butterflies. Genome Biology and Evolution, 10(6), 1351–1362. https://doi.org/10.1093/GBE/EVY093

Trapnell, C., Roberts, A., Goff, L., Pertea, G., Kim, D., Kelley, D. R., Pimentel, H., Salzberg, S. L., Rinn, J. L., & Pachter, L. (2012). Differential gene and transcript expression analysis of RNA-seq experiments with TopHat and Cufflinks. Nature Protocols, 7(3), Article 3. https://doi.org/10.1038/nprot.2012.016

Wang, H., Shi, Y., Wang, L., Liu, S., Wu, S., Yang, Y., Feyereisen, R., & Wu, Y. (2018). CYP6AE gene cluster knockout in *Helicoverpa armigera* reveals role in detoxification of phytochemicals and insecticides. Nature Communications, 9(1), Article 1. https://doi.org/10.1038/s41467-018-07226-6

Weaver, S., Shank, S. D., Spielman, S. J., Li, M., Muse, S. V., & Kosakovsky Pond, S. L. (2018). Datamonkey 2.0: A modern web application for characterizing selective and other evolutionary processes. Molecular Biology and Evolution, 35(3), 773–777. https://doi.org/10.1093/molbev/msx335

Wheat, C. W., Vogel, H., Wittstock, U., Braby, M. F., Underwood, D., & Mitchell-Olds, T. (2007). The genetic basis of a plant–insect coevolutionary key innovation. Proceedings of the National Academy of Sciences, 104(51), 20427–20431. https://doi.org/10.1073/pnas.0706229104

Wiens, J. J., Lapoint, R. T., & Whiteman, N. K. (2015). Herbivory increases diversification across insect clades. Nature Communications, 6(1), Article 1. https://doi.org/10.1038/ncomms9370

Wittstock, U., & Halkier, B. A. (2002). Glucosinolate research in the Arabidopsis era. Trends in Plant Science, 7(6), 263–270. https://doi.org/10.1016/S1360-1385(02)02273-2

You, Y., Xie, M., Ren, N., Cheng, X., Li, J., Ma, X., Zou, M., Vasseur, L., Gurr, G. M., & You, M. (2015). Characterization and expression profiling of glutathione S-transferases in the diamondback moth, *Plutella xylostella* (L.). BMC Genomics, 16(1), 152. https://doi.org/10.1186/s12864-015-1343-5

